# Somatostatin receptor subtypes 1 and 4 redundantly regulate neprilysin, the major amyloid-beta degrading enzyme in brain

**DOI:** 10.1101/2020.05.09.085795

**Authors:** Per Nilsson, Karin Sörgjerd, Naomasa Kakiya, Hiroki Sasaguri, Naoto Watamura, Makoto Shimozawa, Satoshi Tsubuki, Zhulin Zhou, Raul Loera-Valencia, Risa Takamura, Misaki Sekiguchi, Aline Petrich, Stefan Schulz, Takashi Saito, Bengt Winblad, Takaomi C. Saido

## Abstract

Alzheimer’s disease (AD) brains are characterized by increased levels of the pathogenic amyloid beta (Aβ) peptide, which accumulates into extracellular plaques. Finding a way to lower Aβ levels is fundamental for the prevention and treatment of AD. Neprilysin is the major Aβ degrading enzyme which is regulated by the neuropeptide somatostatin. Here we used a combination of *in vitro* and *in vivo* approaches to identify the subtype specificity of the five somatostatin receptors (SSTs) expressed in the brain, involved in the regulation of neprilysin. Using a battery of *Sst* double knockout (dKO) mice we show that neprilysin is regulated by SST_1_ and SST_4_ in a redundant manner. *Sst*_*1*_ and *Sst_4_* dKO mice exhibit a specific decrease of presynaptic neprilysin in the Lacunosum molecular layer. Moreover, a genetic deficiency of *Sst*_1_ and *Sst_4_* in amyloid beta precursor protein (*App*) knock-in mice, an AD mouse model, aggravates the Aβ pathology in the hippocampus. As a first proof of concept towards an Aβ-lowering strategy involving neprilysin, we demonstrate that treatment with an agonist selective for SST_1_ and SST_4_ ameliorates the Aβ pathology and improves cognition in the *App* knock-in AD mouse model.

## Introduction

Amyloid β peptide (Aβ) is central to the development of Alzheimer’s disease (AD) and possibly drives the pathology (Hardy and Selkoe, 2002). Increased levels of Aβ lead to the deposition of Aβ in extracellular plaques, which are accompanied by intracellular neurofibrillary tangles composed of hyperphosphorylated protein tau in AD brains (Alafuzoff et al., 2008; Alafuzoff et al., 2009). Mutations in amyloid beta precursor protein (APP) and APP-processing presenilin genes cause early onset familial AD by increasing total Aβ generation or by specifically increasing the production of aggregation-prone Aβ42 or Aβ43 (Mullan et al., 1992; Welander et al., 2009; Saito et al., 2011). The recent finding that a mutation in APP is protective against AD further strengthens the genetic link between Aβ and AD (Jonsson et al., 2012). In sporadic AD (SAD), which comprises 99% of all AD cases, a strong genetic link has not been identified and a decreased Aβ catabolism may instead underlie the increased Aβ levels. This could be caused by a decrease in the major Aβ-degrading enzyme neprilysin, a membrane-bound metallo-endopeptidase that efficiently degrades Aβ *in vitro* and *in vivo* (Iwata et al., 2000; Iwata et al., 2001). Genetic deletion of neprilysin increases Aβ pathology in a gene dose-dependent manner and worsens memory function in AD mouse models (Huang et al., 2006; Hüttenrauch et al., 2015). Levels of neprilysin in the brain decrease with age and in AD, which could result in an increase of Aβ levels in the brain (Yasojima et al., 2001; Carpentier et al., 2002; Iwata et al., 2002; Maruyama et al., 2005; Hellström-Lindahl et al., 2008). Increasing neprilysin activity may therefore represent a potential Aβ lowering treatment. Indeed, expression of neprilysin by gene therapeutic approaches in mice lowers Aβ pathology and improves cognition (Iwata et al., 2004; Iwata et al., 2013).

The activity of neprilysin is controlled by its translocation to the cell membranes from intracellular vesicles (Saito et al., 2005), which in turn is regulated by its phosphorylation/dephosphorylation (Kakiya et al., 2012). We previously identified in an *in vitro* screening procedure the neuropeptide somatostatin as an activator of neprilysin (Saito et al., 2005). Somatostatin, also known as somatotropin-release inhibiting factor (SRIF), inhibits the release of thyroid stimulating hormone and growth hormone from the pituitary gland. SRIF also plays a role in the gastrointestinal tract where it inhibits the secretion of insulin and secretin, and controls intestinal motility (Corleto, 2010). SRIF levels decrease in the brain with aging and in AD, possibly due to the degeneration of SRIF-expressing interneurons (Davies et al., 1980; Beal et al., 1985; Bergström et al., 1991; Hayashi et al., 1997; van de Nes et al., 2002; Lu et al., 2004; Gahete et al., 2010). In addition, single nucleotide polymorphisms (SNPs) in the SRIF gene have been linked to AD (Vepsäläinen et al., 2007). SRIF signalling is mediated by five somatostatin receptor (SST) subtypes, which are G-protein coupled receptors (GPCR). To study the Aβ pathology related to AD in mice, novel *App* knock-in mouse models have been developed (Saito et al., 2014). These models are free of APP overexpression and thus artefacts associated with the high and unphysiological levels of APP in APP transgenic mice can be circumvented (Saito et al., 2014; Sasaguri et al., 2017).

*App*^*NL-F*^ knock-in mice harbour the Swedish and the Beyreuther/Iberian mutations which induce high Aβ42 levels as well as a high Aβ42/Aβ40 ratio, as seen in AD brains. These mice develop Aβ plaques from nine months of age and cognitive impairments from 15 months of age (Saito et al., 2014). In addition to the Swedish and the Beyreuther/Iberian mutations, *App*^*NL-G-F*^ knock-in mice harbour the arctic mutation which enhances Aβ oligomerization properties (Nilsberth et al., 2001). These mice develop Aβ plaques from two months of age and memory deficits from six months of age. Having a different age of onset and progress of pathological properties, the two mouse models can thus serve different research purposes. The rapid development of AD pathology in *App*^*NL-G-F*^ mice offers the possibility for a shorter treatment time, while the long-term effects of gene manipulation can be studied in *App*^*NL-F*^ mice, which exhibit a slower progression of AD-like pathologies.

Previous studies have indicated the involvement of SST_4_ in the regulation of neprilysin. This was shown by using either *i.p.* or *i.c.v.* injection of the SST_4_ agonist NNC 26-9100 to wildtype and SAMP8 mice (Sandoval et al., 2011; Sandoval et al., 2012, 2013). However, *in vivo* data obtained from genetically modified mice on the SRIF-induced regulation of neprilysin have been missing. Here, we have used a battery of *Sst* subtype knockout (KO) mice and show that SST_1_ and SST_4_ subtypes redundantly regulate neprilysin. As a first step towards developing a treatment to lower Aβ levels in the brain through SRIF-induced neprilysin activation, we also show that the Aβ pathology in *App*-knock-in mice can be ameliorated by delivery of an agonist selective to SST_1_ and SST_4_.

## Results

We previously developed an *in vitro* system that detects neprilysin activity in living cells (Saito et al., 2005). To achieve a robust SRIF-induced activation of neprilysin in this primary neuron-based cell culture system, a mixture of wildtype hippocampal, cortical and striatal neurons (in a 9:9:1 ratio) is required. By adding SST subtype-specific agonists to this *in vitro* system, we found that SST_1_ and SST_4_-specific agonists activated neprilysin, whereas co-treatment with cyclo-SRIF, a competitive antagonist, abolished this activation (Fig 1a). This finding indicates the involvement of SST_1_ and SST_4_ in the activation of neprilysin in *vitro*. Indeed, primary neurons derived from *Sst_1_xSst_4_* dKO mice exhibited lower neprilysin activity while primary neurons from *Sst*_*1*_ and *Sst_4_* KO mice exhibited unaltered neprilysin activity (Fig 1b). This led to increased Aβ levels in the conditioned media of the cell cultures, without detectable changes in cell viability of the *Sst_1_xSst _4_* dKO neurons, confirming a role for SST_1_ and SST_4_ in neprilysin regulation (Fig 1c,d). The involvement of SST_1_ and SST_4_ in the regulation of neprilysin was also confirmed using our previously established neprilysin activity staining method that visualizes neprilysin located on the cell surface (Saito et al., 2005). The addition of SRIF increased neprilysin in wildtype neurons but did not activate neprilysin in neurons from *Sst_1_xSst_4_* dKO mice, which exhibited lower neprilysin activity as compared to wildtype neurons (Fig 2a, b).

**Figure 1.**
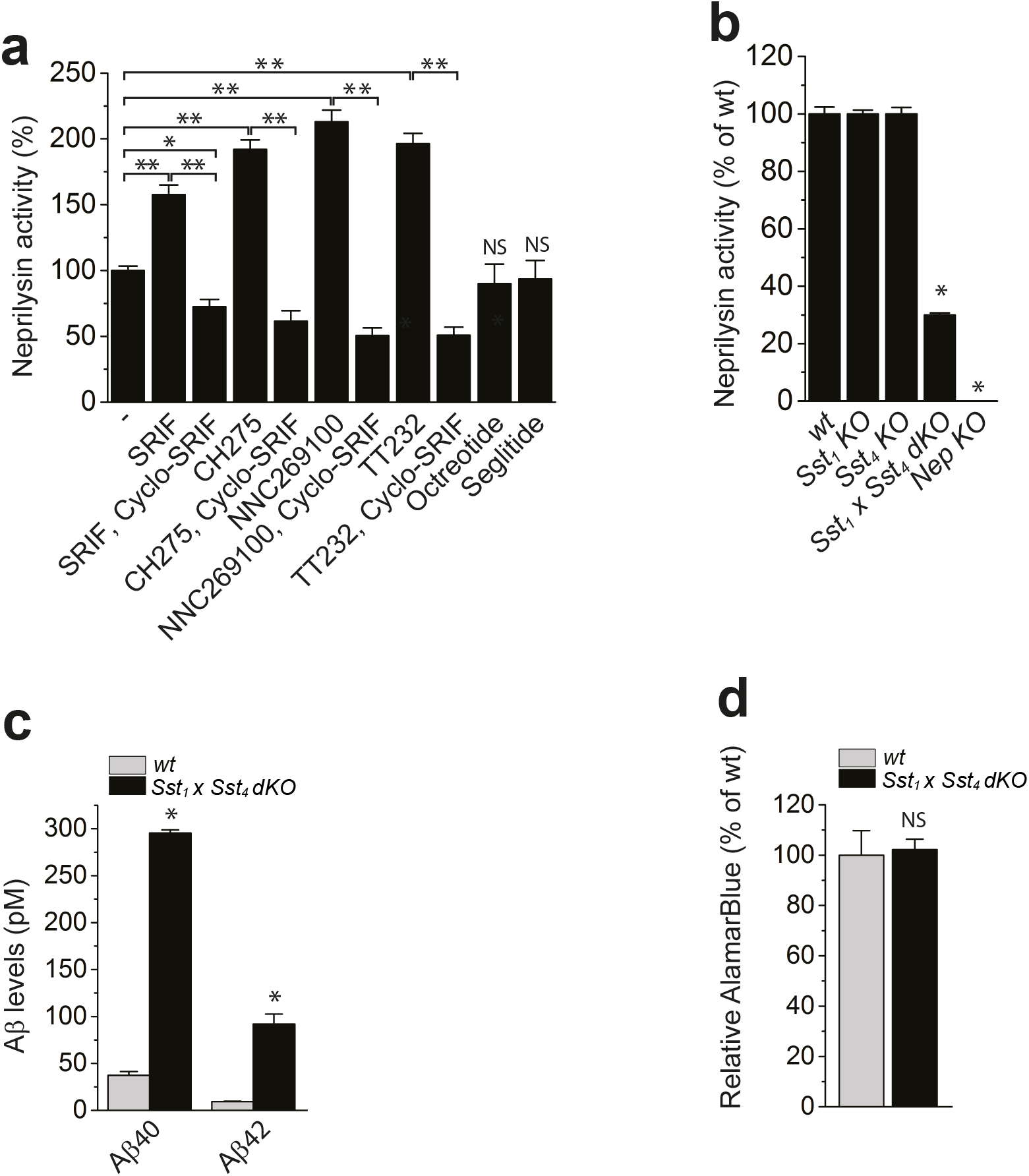
SST_1_ and SST_4_ regulate neprilysin and Aβ levels *in vitro*. (a) Neprilysin activity in wildtype primary neurons exposed to somatostatin (SRIF), CH275 (SST_1_ + SST_4_ agonist), NNC269100 (SST_4_ agonist), TT232 (SST_1_ + SST_4_ agonist), Octreotide (SST_2_, SST_3_ and SST_5_ agonist) and Seglitide (SST_2_, SST_3_ and SST_5_ agonist) either alone or in combination with cyclo-SRIF. n = 3-4 per treatment, \**p* < 0.005, \*\**p* < 0.001. (b) Neprilysin activity in primary neurons derived from mice with indicated genotypes. n = 12/condition and repeated with at least 3 different batches of neurons from 3 different animals. \**p* < 0.01. (c) Aβ levels in conditioned media from primary neurons derived from wildtype and *Sst_1_xSst_4_* dKO mice. n = 4, \**p* < 0.005. (d) AlamarBlue staining of the conditioned media from wildtype and *Sst_1_xSst_4_* dKO primary neurons. n = 4, no significant difference.

**Figure 2.**
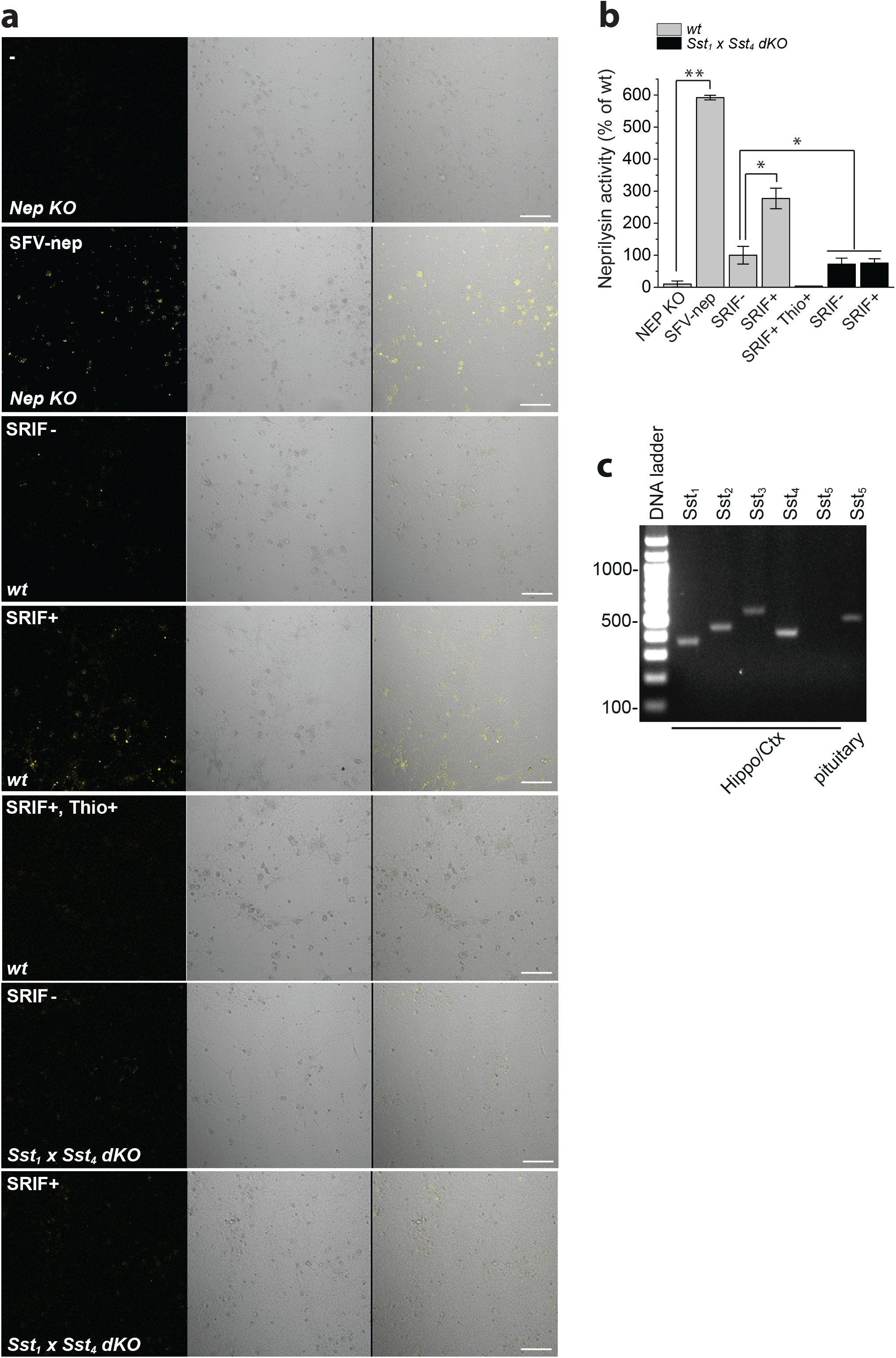
*In vitro* activity staining of neprilysin in primary neurons. (a) Staining of neprilysin activity to visualize cell surface-located neprilysin in primary neurons obtained from mice with genotypes as indicated with and treated as indicated were quantified. (b) The fluorescence signal was quantified. n = 3, \**p* < 0.05, \*\**p* < 0.001. (c) RT-PCR analysis of SST subtype expression in cortex/hippocampus. SFV; semlikiforest virus, SRIF; somatostatin, Thio; thiorphan

In agreement with a previous study (Schulz et al., 2000), we next confirmed that SST_1_, SST_2_, SST_3_ and SST_4_ are expressed in the hippocampus and cortex (regions primarily affected by AD pathology), whereas SST_5_ is exclusively expressed in the pituitary (Fig 2c). We therefore continued to analyse the *in vivo* effects of *Sst_1_-Sst_4_* genetic deficiency in the brain of the KO mice. Whereas neprilysin levels were not altered in 3-month-old single *Sst* KO mice, *Sst_1_xSst_4_* dKO mice exhibited a significant decrease of neprilysin specifically in the Lacunosum molecular layer (Lmol) where SRIF is highly expressed (Fig 3 a, b, c, 4a). No other *Sst* dKO combinations led to changes in neprilysin levels. A decrease of neprilysin in the Lmol layer was also observed in *Srif* KO mice (Fig 4a). Consistent with the immunohistochemical data, a significant decrease in neprilysin levels in hippocampal brain homogenates from *Sst_1_xSst_4_* dKO mice was confirmed by a neprilysin-specific ELISA (Fig 3d). Furthermore, the specific decrease of neprilysin upon deletion of *Sst*_*1*_ and *Sst_4_* caused Aβ42 levels to increase in hippocampus of the *Sst_1_*x*Sst_4_* dKO mice, in contrast to the unaltered Aβ levels in single *Sst*_*1*_ or *Sst_4_* KO mice (Fig 3e, f and data not shown). The significantly increased and accumulated Aβ levels may have contributed to the detected impaired memory in the *Sst_1_xSst_4_* dKO mice (Fig 3g,h). However, the impairment could also be due to other SST-mediated effects caused by the deficiency of *Sst*_*1*_ and *Sst_4_*. To directly visualize the impact of *Sst*_*1*_ and *Sst_4_* deficiency on Aβ pathology, we crossed the *Sst_1_xSst_4_* dKO mice with an *App* knock-in AD mouse model, *App*^*NL-F*^ (Saito et al., 2014), which begins to exhibit Aβ plaques from nine months of age. Strikingly, the Aβ pathology in the Lmol layer of hippocampus of 15-month-old *Sst_1_xSst_4_ dKO x App^NL-F^* mice was significantly enhanced, as shown by an increase in both the number and size of Aβ plaques, where presynaptic neprilysin levels, as visualized with co-immunostaining for neprilysin and the presynaptic marker synaptophysin, were decreased upon deletion of *Sst*_*1*_ and *Sst_4_* (Fig 4a, b, c, d). In addition, we confirmed that genetic deletion of *Sst*_*1*_ and *Sst_4_* did not affect APP processing nor the levels of another Aβ-degrading enzyme, insulin degrading enzyme (IDE), (Fig 4e, f).

**Figure 3.**
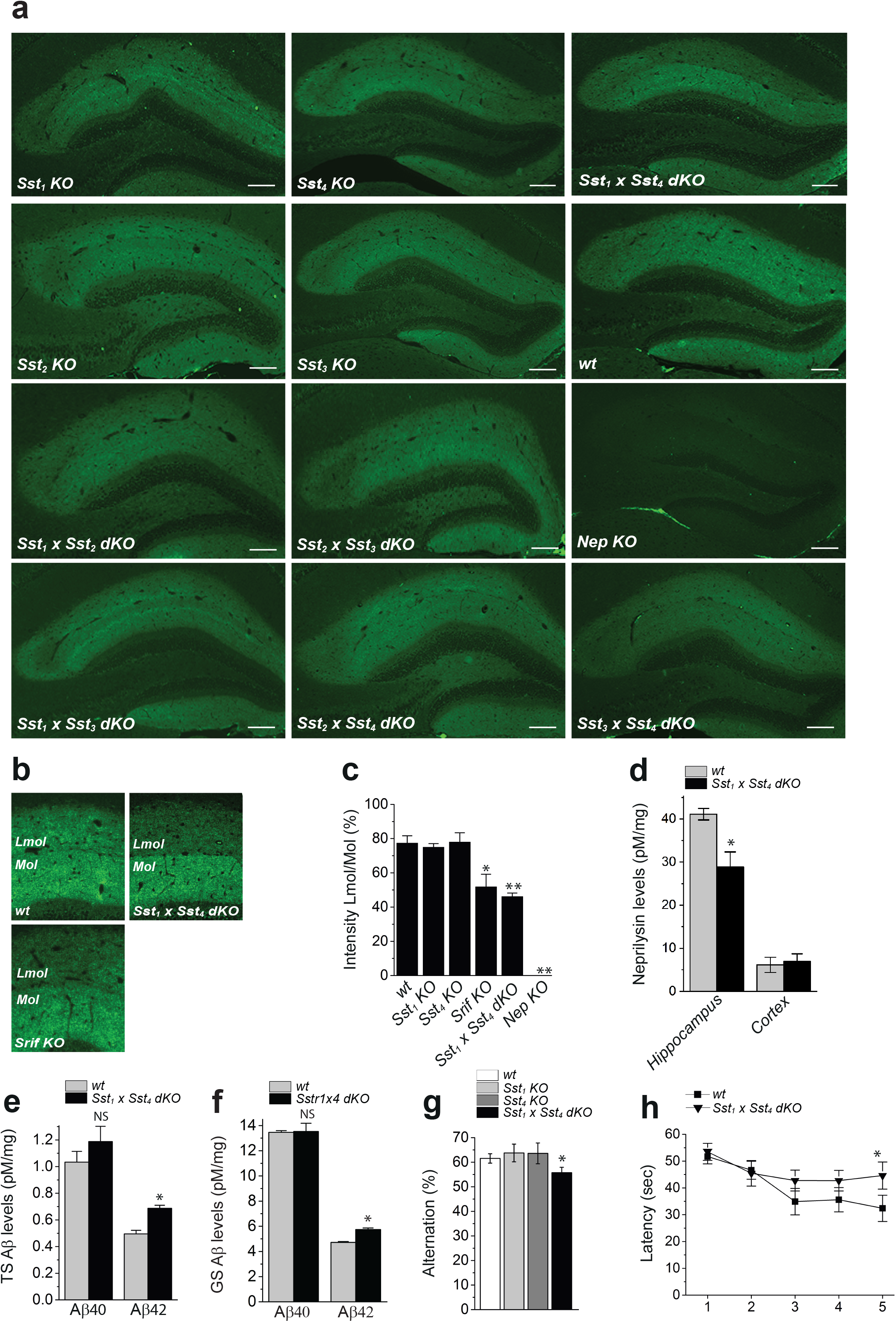
*Sst*_*1*_ and *Sst*_*4*_ genetic deficiency leads to decreased neprilysin in Lacunosum molecular layer of hippocampus. (a) Genetic deletion of SST_1_ and SST_4_ specifically decreases neprilysin in Lmol layer of hippocampus. Neprilysin immunostaining in the hippocampus of brain sections from three months old mice with genotypes as indicated. (b) Neprilysin staining in the zoomed-in areas of the molecular (Mol) and Lacunosum molecular (Lmol) layers was quantified (c). n = 3-4, \**p* < 0.05 and \*\**p* < 0.005. (d) Neprilysin levels in the hippocampus of wildtype and *Sst_1_xSst_4_* dKO mice were determined by neprilysin ELISA. n = 3-4, \**p* < 0.05. (e) Tris-soluble (TS) and (f) Gu-HCl-soluble Aβ levels in the hippocampus were determined by Aβ ELISA. n = 3, \**p* < 0.0005. Memory test of mice by Y-maze (g) and Morris water maze (h). n = 15, \**p* < 0.05. Scale bars in (e): 100 μm.

**Figure 4.**
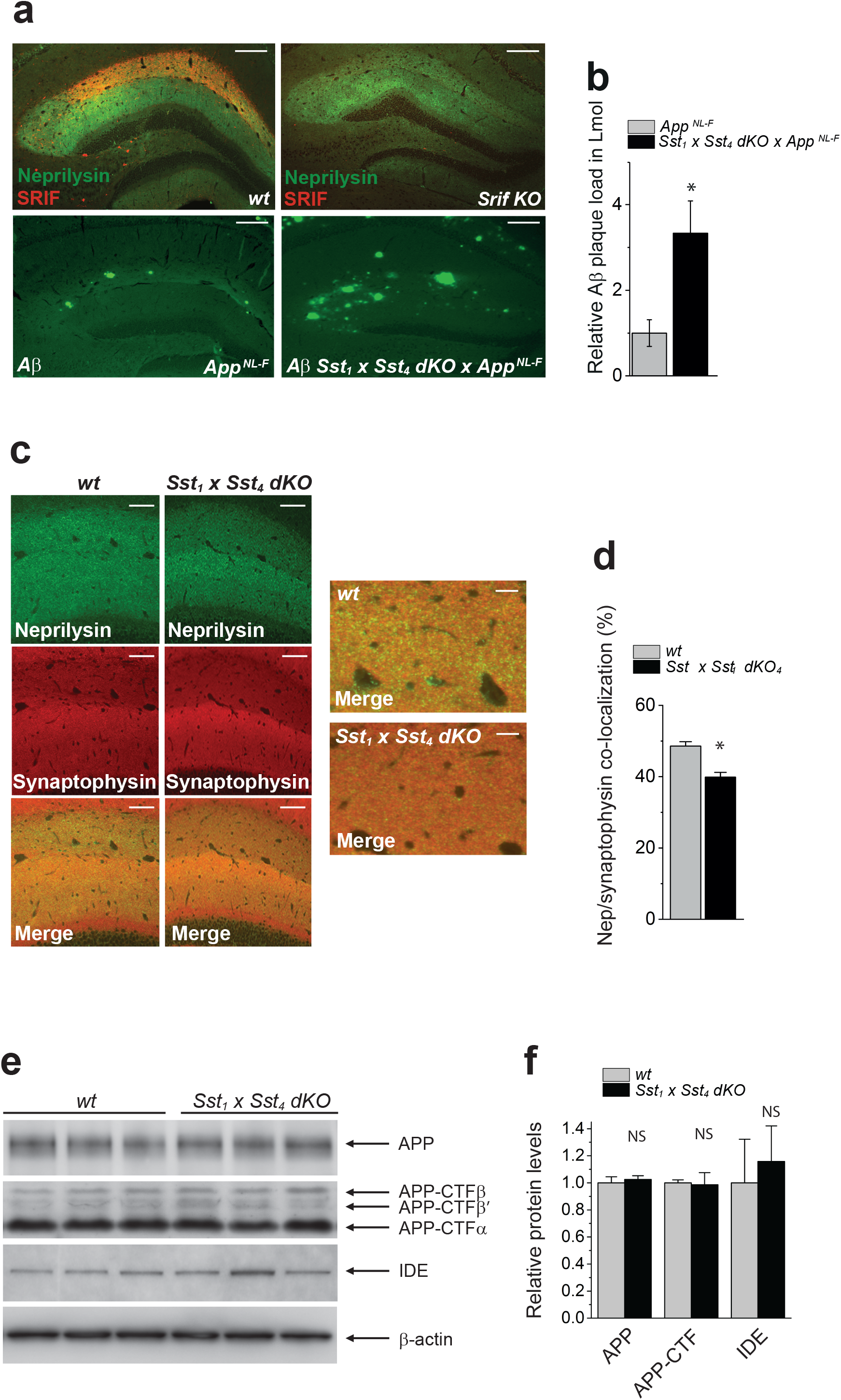
A dual deficiency of *Sst*_*1*_ and Sst_*4*_ increases Aβ pathology in single *App* knock-in mice. (a) A lack of SST_1_ and SST_4_ enhances the Aβ pathology in *App*^*NL-F*^ knock-in mice. Double staining for neprilysin and SRIF of wildtype and *Srif* KO mouse brain (upper two panels). In the lower two panels, 15 months old *App*^*NL-F*^ and *Sst_1_xSst_4_* dKO x *App*^*NL-F*^ mice were stained for Aβ. (b) Aβ plaque load in Lmol layer was quantified and presented as relative plaque load comparing *App*^*NL-F*^ to *Sst_1_xSst_4_* dKO x *App*^*NL-F*^ mice. n = 3, \**p* < 0.005. (c) Co-immunostaining for neprilysin and synaptophysin in wildtype and *Sst_1_xSst_4_* dKO mice, in the zoomed-in panels a region of Lmol layer is shown. (d) Co-localization was quantified. n = 3-4, \**p* < 0.05. (e, f) APP, APP-CTF and insulin-degrading enzyme (IDE) levels in the hippocampus of wildtype and *Sst_1_xSst_4_* dKO mice were analyzed by quantitative western blot. β-actin was used as a loading control. No significant differences were observed, n = 3. Scale bars is 100 μm in (a) and (c) left panels and 30 μm in (c) right panels.

To further investigate the effect of ageing on the SRIF-neprilysin regulatory system, 25-month-old *App*^*NL-F*^ mice were examined and compared to 3-month-old *App*^*NL-F*^ mice. Interestingly, the levels of both SRIF and neprilysin in Lmol and Mol layer decreased upon ageing as similarly observed in aged TgCRND8 mice (Fukami et al., 2002) (Fig 5a, b, c, d). This is paralleled by dense and area-specific Aβ deposits in the molecular layer and dentate gyrus. However, Aβ plaque load was significantly lower in Lmol layer as compared to Mol layer (Fig 5e).

**Figure 5.**
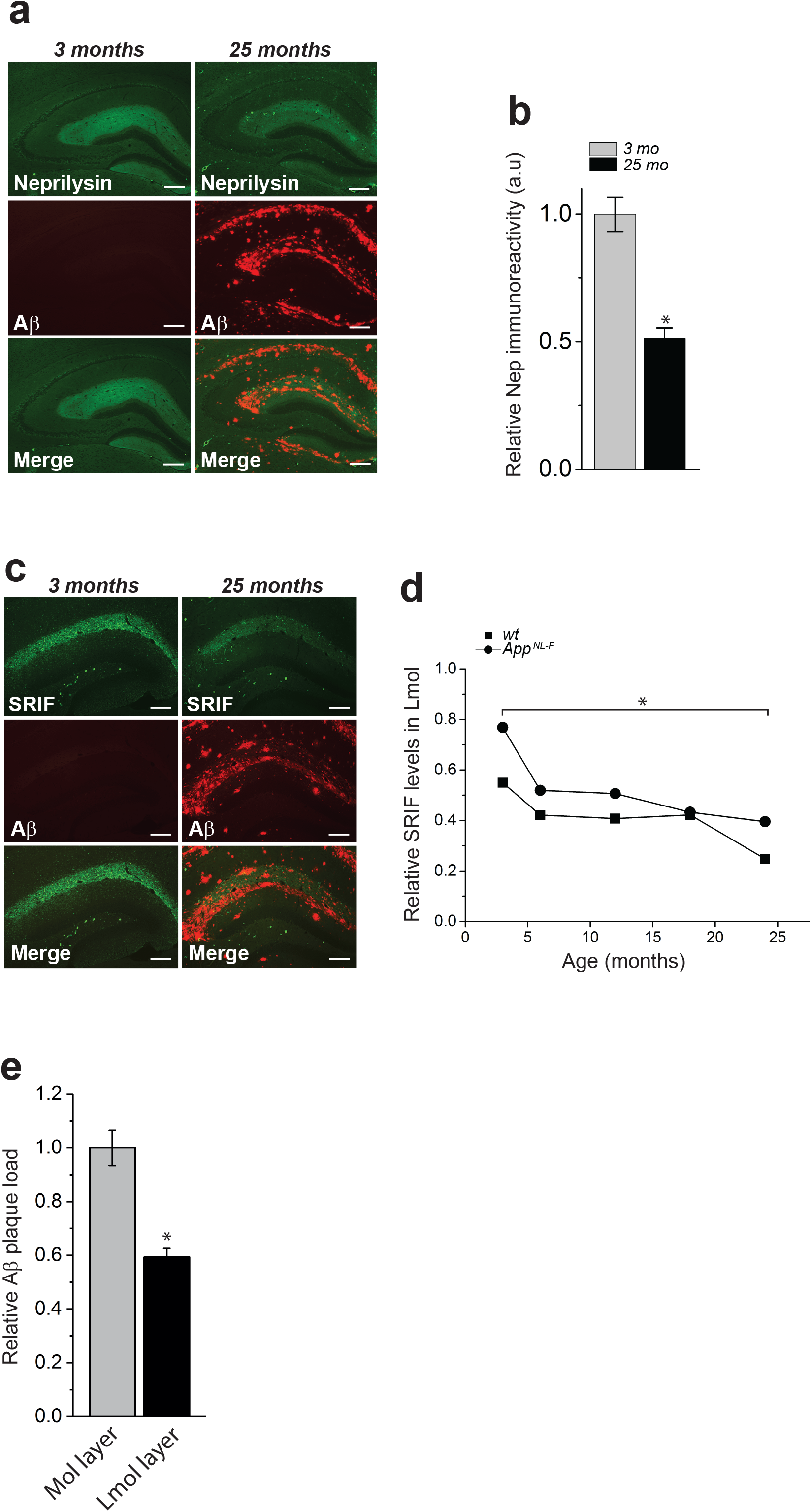
Age-related decrease of neprilysin and somatostatin in *App*^*NL-F*^ knock-in mice. (a,b) Co-immunostaining of neprilysin and Aβ revealed lowered neprilysin levels with increased age; this was associated with Aβ deposits in the molecular and Lmol layers. n = 3-4, \**p* < 0.005 (c,d) Co-immunostaining of somatostatin (SRIF) and Aβ demonstrated decreased SRIF with age in the Lmol layer of *App*^*NL-F*^ knock-in mice, paralleled by Aβ plaque depositions. n = 3-4, \**p* < 0.05 e) Quantification of Aβ plaque load in Mol and Lmol. n = 4, \**p* < 0.01. Scale bars 200 μm.

To address if the decreased levels of neprilysin/SRIF in the *App* knock-in mice can be activated by administration of an SST subtype specific agonist, we treated *App^NL-^G-F* mice which rapidly develop a robust Aβ pathology, with an SST_1_/SST_4_ agonist. Previous studies have indicated that both *i.p* and *i.c.v* injection of the SST_4_ agonist NNC 26-9100 lowers Aβ levels and improves memory in SAMP8 and APP Tg2576 mice (Sandoval et al., 2011; Sandoval et al., 2012, 2013). However, two weeks’ administration of NNC 26-9100 by *i.p.* injections to *App*^*NL-F*^ mice did not lower the Aβ levels (data not shown). We therefore choose to administer the SST_1_ and SST_4_ agonist CH275 since 1) SST_1_ and SST_4_ expression is high in the hippocampus and cortex compared to other vital non-neural organs (Fig 6a, b) and 2) a PBS-soluble SST_1_ and SST_4_ agonist is commercially available. However, the peptide-based CH275 does not cross the blood-brain-barrier (BBB) and we therefore administered CH275 or PBS directly into the Lmol layer of 2-month-old *App*^*NL-G-F*^ mice for four months. *App*^*NL-G-F*^ mice begin to exhibit Aβ plaques at two months of age. This treatment robustly increased the expression of neprilysin in hippocampus which was paralleled by a clear reduction in Aβ plaque load in the same region, as compared to the contralateral side, and without causing any toxic side effects (Fig 6c). A trend towards decreased TS and GS Aβ42 levels was additionally measured by an Aβ-specific ELISA (*p* = 0.1, data not shown).

**Figure 6.**
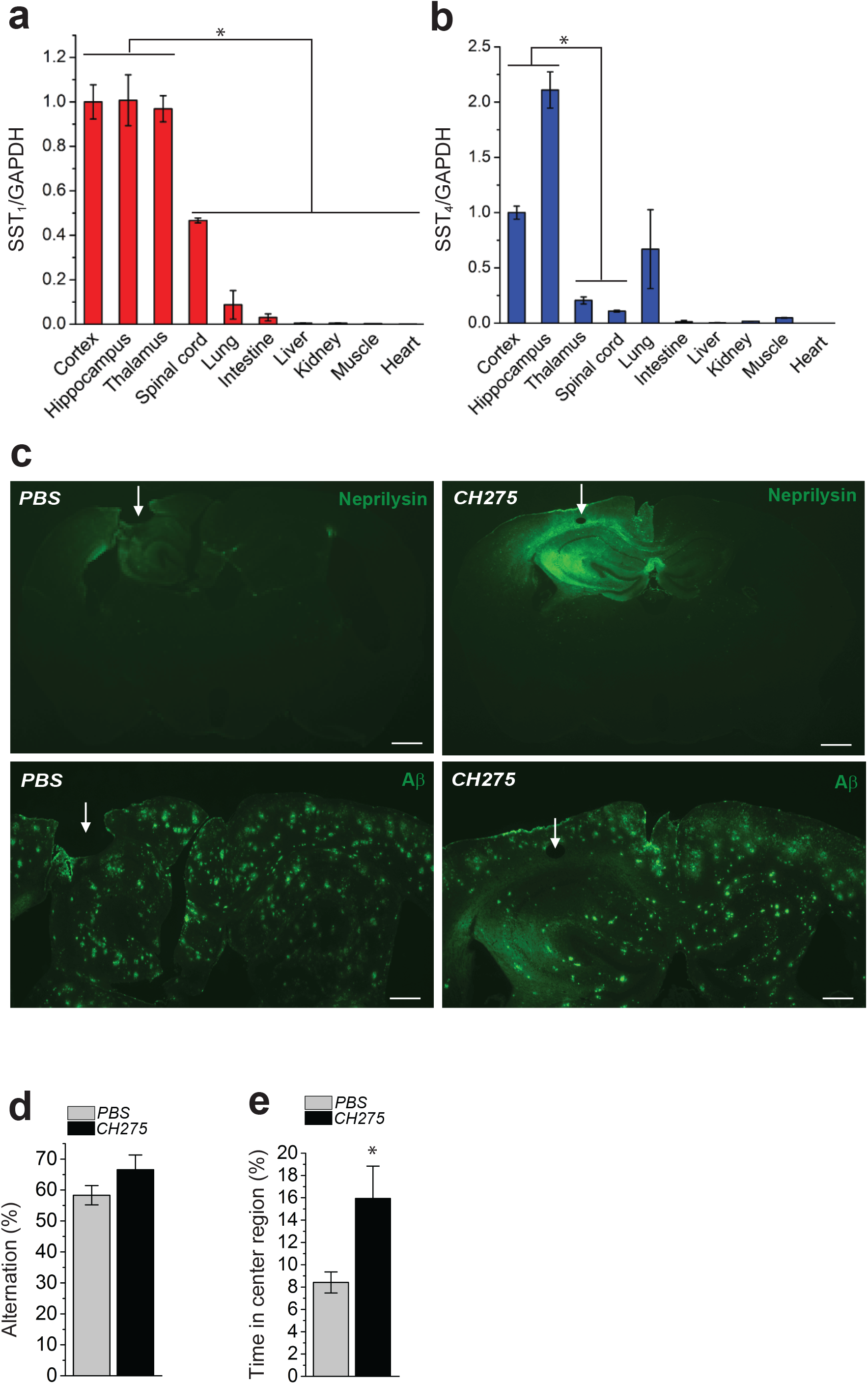
Administration of SST_1_ and SST_4_ agonist increases neprilysin and lowers Aβ levels in *App*^*NL-G-F*^ knock-in mice. (a,b) SST_1_ and SST_4_ expression analysed by qPCR and normalized to GAPDH in tissues as indicated. n = 3, \**p* < 0.005. PBS or CH275 was administered for four months by osmotic pumps implanted in 2-month-old *App*^*NL-G-F*^ mice. (c) Neprilysin and Aβ plaque load in hippocampus of treated mice as shown by immunohistochemical analysis. (d,e) Mouse behaviour after four months of treatment was analysed by Y-maze (d) and open field tests (e), n = 9-11 mice/treatment. Arrows indicate the site of cannula placement. \**p* < 0.05. Bar indicates 500 μm in upper panels and 300 μm in lower panels.

We also observed a tendency towards improved memory performance of the agonist-treated mice in the Y-maze after the administration period (*p* = 0.08), while the open field test revealed a significantly reduced level of anxiety in these mice (*p* = 0.02) (Fig 6d, e). The improved cognition could be due to the decreased Aβ pathology; however, an effect on behaviour of CH275 that potentially remained in the brain cannot be excluded even though the pumps were confirmed empty when removed from the mouse brains. In addition, measurement of the concentration of CH275 in dissected brain hippocampus rendered an approximate total concentration of 2 μM (Fig 7a, b, c). Taken together, these findings demonstrate in a proof of concept that a treatment based on the administration of an SST_1_/SST_4_ agonist(s) could lower Aβ amyloidosis and ameliorate cognitive impairment in a relevant mouse model of AD.

**Figure 7.**
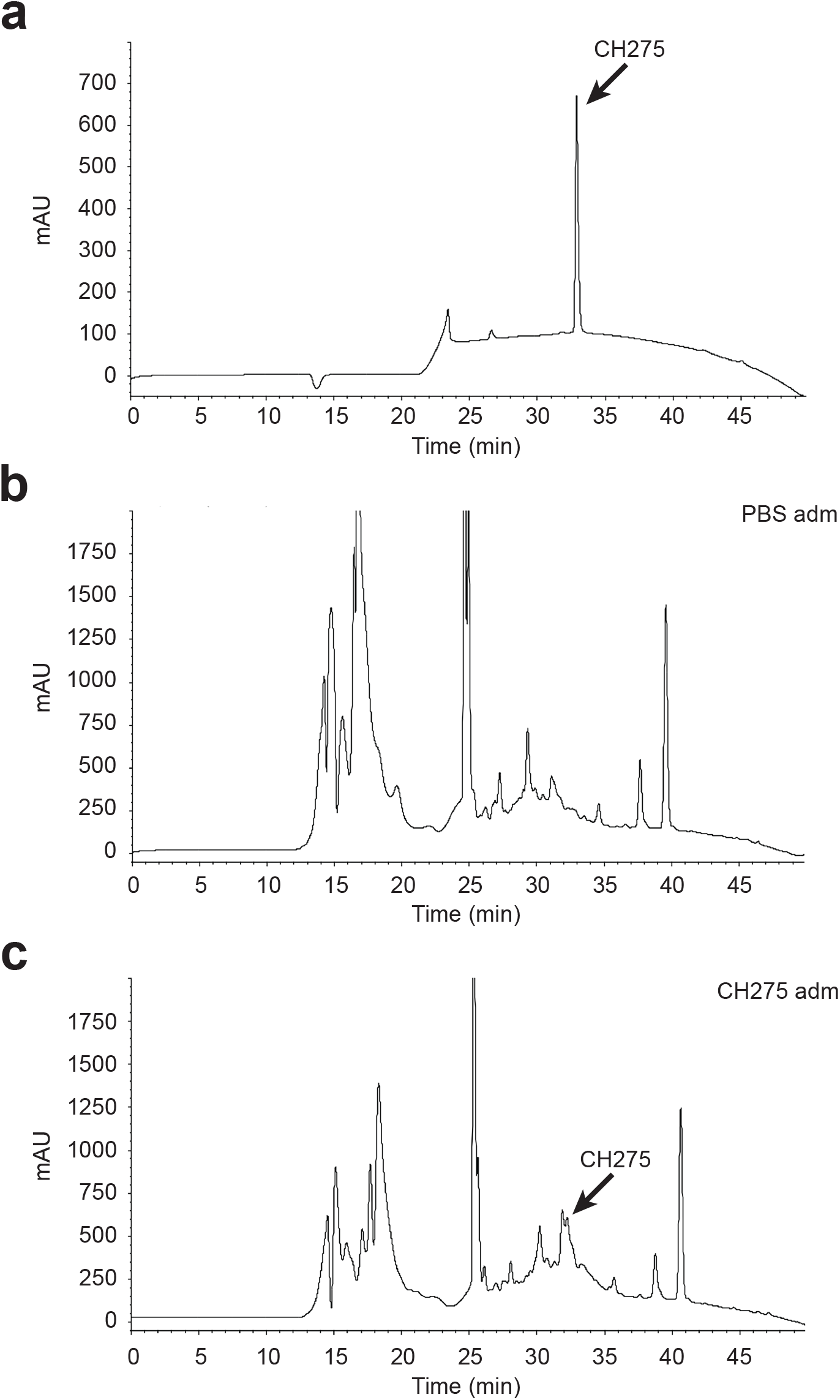
Confirmation of delivery and *in vivo* concentration of SST_1_ and SST_4_ agonist CH275 in hippocampus. (a) 2 μg pure CH275 was analyzed by HPLC and used as standard to measure the area under the peak, 8318.5 mAU × time. (b, c) The homogenates of brains administered with PBS or CH275 were analyzed by reversed phase HPLC. The area under the peak corresponding to CH275 in the CH275-treated brain homogenate was 4349.0 mAU × time. The concentration of CH275 in the brain homogenate was 34.395 ng/mg [(4349.0/8318.5) × 2 μg / 30.4 mg].

## Discussion

We have shown in this report, by using a combination of *in vitro* and *in vivo* paradigms, that SST_1_ and SST_4_, which belong to the same SST family, SRIF2, redundantly regulate the major Aβ degrading enzyme neprilysin. A genetic deletion of both SST_1_ and SST_4_ leads to a specific decrease of neprilysin in Lacunosum molecular layer, a region rich is synapses and where both SRIF and neprilysin is present at high levels. SSTs are known to dimerize and it is possible that SST_1_ and SST_4_ either form functional homodimers or heterodimers, although this remains to be shown biochemically or biophysically. The need to have striatal neurons to facilitate maximum activation of neuronal neprilysin by SRIF *in vitro*, also implies the presence of molecular interaction(s) between cortical/hippocampal and striatal neurons. Although the molecular mechanisms that account for the neprilysin activation remain elusive, phosphorylation and dephosphorylation of the neprilysin cytoplasmic domain by Mitogen-activated Protein Kinase/Extracellular Signal-regulated Kinase Kinase (MEK) and Protein Phosphatase 1a (PP1a), respectively, is likely to play a major role (Kakiya et al., 2012).

We found here that the double genetic deficiency of *Sst*_*1*_ and *Sst*_*4*_ specifically decrease neprilysin in Lmol layer of hippocampus which leads to increased Aβ42 levels and Aβ plaque pathology, which are commonly increased in both SAD and FAD. Thus, it is possible that the decreased levels of SRIF observed in ageing mouse and human brains (Iwata et al., 2002; Hellström-Lindahl et al., 2008) and in AD brains (Davies et al., 1980) lead to decreased neprilysin levels. This would likely result in an accumulation of Aβ over time, which activates pathological events downstream of Aβ and the onset of AD. SST_1_ and SST_4_ together have the most brain-specific expression pattern (Fig 6), meaning that a therapeutic strategy with BBB-permeable SST_1_ and/or SST_4_ agonists could be beneficial to slow the progression of AD. Such a treatment, possibly in combination with other disease-modifying medications targeting, *e.g*., the tau pathologies, may also reduce anxiety, a prominent symptom of AD, since SST agonist are known to reduce anxiety. Given that more than 400 medication candidates for AD, including immunotherapies, have failed in clinical trials, now may be the time to consider low-molecular-weight medication(s) that target G protein-coupled receptors (GPCRs), given that ligand-receptor binding in GPCR(s) is one of the most specific and selective interactions in biological systems, generating minimal side effects. In addition, such synthetic compounds should be cheaper to produce than immunotherapies and would thus be highly valued in an increasingly aging society.

Specific indications from these data in this research include that the following may become the preventive/therapeutic targets as “specific GPCR(s)” via regulation of Aβ metabolism by 1) SST_1_ homodimer 2) SST_4_ homodimer 3) SST_1_ homodimer and SST_4_ homodimer 4) SST_1_ and SST_4_ heterodimer(s) (Kossut et al., 2018). Identification of the relevant target molecule among these will be important for generation of specific and elective medication(s), but it is beyond the scope of this research. Finally, the prolonged elevation of neprilysin expression upon CH275 administration indicates that the target SST is likely free from receptor desensitization, which would affect its medical utility.

## Acknowledgments

We express our gratitude to members of RIKEN PNS laboratory. We thank Ute Hochgeschwender, Oklahoma Medical Research Foundation, for providing *Srif KO*, *Sst_3_ KO* and *Sst_4_ KO* mice, and Craig Gerard, Harvard Medical School for providing the *neprilysin KO* mice. This research was supported financially by grants from the Swedish Research Council, Sweden (PN, BW); Hållsten Research Foundation, Sweden (PN); Swedish Alzheimer Foundation, Sweden (PN); RIKEN Brain Science Institute, Ministry of Education, Sports, Science and Technology, Japan; and the Ministry of Health, Labour and Welfare, Japan (TCS). Support in part was also provided by the Strategic Research Program for Brain Sciences from the Japan Agency for Medical Research and Development, AMED (TS).

## Materials and Methods

### Animals

*Sst*_*1*_ KO mice were obtained from Jackson laboratories; *Sst*_*2*_ KO mice were a kind gift from Stefan Schulz, Friedrich-Schiller University, Germany; *Srif* KO, *Sst*_*3*_ KO and *Sst_4_* KO mice were kindly provided by Ute Hochgeschwender, Oklahoma Medical Research Foundation, USA; and *neprilysin* KO mice were kindly provided by Craig Gerard, Harvard Medical School, USA. Details of the *App*^*NL-F*^ mice have been previously published^17^. All mice were bred on a C57BL/6 background. The animal experiments were conducted according to the guidelines of the RIKEN Center for Brain Science.

### Primary cultures

Primary neuronal co-cultures consisting of cortical, hippocampal and striatal neurons in a ratio of 9:9:1, which is necessary to stably and reproducibly recapitulate the SRIF stimulatory effect in vitro, were prepared by dissecting the respective brains regions from E17-18 C57BL/6J, *Sst*_*1*_ KO, *Sst_4_* KO, *Sst_1_xSst_4_* dKO and *neprilysin* KO mouse embryos, and plated in 96-well plates at a total density of 2.5 × 10^4^ cells. For neprilysin activity staining, 1.8 × 10^5^ viable cells were plated in 24-well plates with poly-L-Lysine-coated coverslips.

### Neprilysin ELISA

Membrane preparations of hippocampal brain homogenates from wildtype and *Sst_1_xSst_4_* dKO mice were prepared by homogenizing the dissected tissue in ice cold 10 mM Tris, pH 8.0, 0.25 M sucrose and EDTA-free protease inhibitors (Roche, cat# 05056489001) followed by centrifugation at 9000 rpm for 15 minutes. The supernatant was recovered and centrifuged at 30,000 rpm for 30 min. The resulting pellet was dissolved in 240 μl 10 mM Tris, pH 8.0, 0.25 M sucrose, EDTA-free protease inhibitors and 1 % Triton-X for 1 hour at 4°C. Solubilized material was centrifuged at 70,000 rpm for 20 minutes, the protein concentration was determined by BCA (Pierce, cat# 23225), and equal amounts of protein were used in the neprilysin ELISA (R&D, cat# DY1126), which was processed according to the manufacturer’s instructions.

### Aβ ELISA

Tris-soluble and guanidine-soluble Aβ levels from hippocampal brain homogenates, prepared as previously described (Saito et al., 2011). Aβ levels in conditioned media was measured by ELISA after addition of guanidine-HCl to the conditioned media to a final concentration of 0.5 M (Aβ40, Wako #294-64701, Aβ42 Wako #290-62601) according to the manufacturer’s instructions.

### Neprilysin activity assay

Neprilysin activity measurements were performed on primary neurons after 24 days *in vitro* (DIV24), without changing the media, as previously described (Kakiya et al., 2012). After a 24-h treatment with 1 μM SRIF (Peptide Institute #4023), 1 μM cyclo-SST (Tocris #3493), 100 nM CH275 (Tocris #2454), 100 nM NNC269100 (Tocris #2440), 1 μM TT232 (Tocris, #4639), 10 nM Octreotide (Sigma, #O1014) or Seglitide (Sigma, #S1316), the cells were incubated with substrate mixture consisting of 50 μM suc-Ala-Ala-Phe-MCA (Sigma, cat# S8758), 1 μM benzyloxycarbonyl (Z)-Leu-Leu-Leucinal (protease inhibitor, synthesized in-house) and Complete EDTA-free Protease Inhibitor (Roche, cat# 05056489001) in 50 mm MES pH 6.5 with or without 10 μM thiorphan (neprilysin-specific inhibitor, Sigma, cat# T6031) at 37 °C for 30 min. Next, 0.1 mg/ml leucine aminopeptidase (Sigma, cat# L-5006) and 0.1 mM phosphoramidon (Peptide institute, cat# 4082) were added, and the reaction mixture was incubated at 37 °C for a further 30 min. 7-Amino-4-methylcoumarin fluorescence was measured at excitation and emission wavelengths of 380 and 460 nm, respectively.

### Neprilysin activity staining

Neprilysin activity staining was performed on DIV24 primary neurons, without changing the media, as previously described (Kakiya et al., 2012). As a control, primary neurons from neprilysin-deficient mice were stained with or without expression of neprilysin by infection with a semliki-forest virus containing the *Mme* gene. After incubation with 1 μM SRIF for 24 hours, the cells were washed once with TBS, fixed in 1.5% PFA, washed twice with TBS and incubated with 200 μl substrate mixture consisting of 0.25 mM glutaryl-Ala-Ala-Phe-methoxy-2-naphtylamide (Sigma, cat# G3769) and 62.5 mM Mes pH 6.8 with or without 10 μM thiorphan for 2 hours at 37 °C. Leucine aminopeptidase (Sigma, #L-5006), phosphoramidon (Peptide Institute, cat# 4082) and 2-hydroxy-5-nitrobenzaldehyde (Acros Organics, #416180050 dissolved in ethanol to 120 mM and centrifuged for one minute at 15,000 rpm to remove insoluble material) were then added to the substrate solution to a final concentration of 10 μg/ml, 20 μM, and 12 mM, respectively, and incubated for 30 min at 37 °C. The cells were washed three times with ice-cold 50 mM Tris pH 7.4, and the cover slips were mounted. Cleaved substrate was visualized by immunofluorescence with a rhodamine filter and argon laser excitation.

### Immunohistochemistry

Four micrometre-thick paraffin-embedded brain sections were autoclaved for 5 minutes for antigen retrieval, immunostained with antibodies against neprilysin (56C6, Novocastra cat# NCL-L-CD10-270), synaptophysin (SY38, ProGen, cat# 61412), SRIF (Millipore, cat# MAB354) and Aβ (82E1, IBL, cat# 10323). Quantification was performed with MetaMorph imaging software (Universal Imaging). Image processing of synaptophysin/neprilysin co-staining was carried out using the FIJI package for Image J (version 1.52p). Images were analyzed at 8-bit grayscale format. Intensity saturation for all images was corroborated using the histogram-analysis function and confirming that no pixels were left on the 255-bin portion of the histogram logarithmic scale. Colocalization between channels in the tissues was carried out using the Coloc2 plugin for FIJI. The signal from the synaptophysin and neprilyisn channels were automatically thresholded. Neprilysin signal was used to create a binary mask that was applied to the synaptophysin signal and the resulting area stained was calculated as percentage of the whole image, equivalent to the degree of colocalization between the two signals.

### Immunoblot analysis

A total of 2.5 μg of Triton-X (1%)-soluble brain homogenates were separated by SDS-PAGE, transferred to PVDF membranes and probed with anti-APP antibody 6E10 (Covance # SIG-39320), anti-β-actin antibody (Sigma, cat# A5441) and anti-IDE (Abcam, cat# ab32216). A total of 25 μg delipidated brain homogenate was separated by SDS-PAGE, transferred to nitrocellulose membrane and probed with anti-APP-CTF antibody (Sigma, cat# A8717).

### RT-PCR

Total RNA was isolated from dissected snap frozen cortical/hippocampal brain tissue and pituitary by RNAiso Plus (Takara cat# 9108). 0.5 ng RNA was reverse transcribed and PCR amplified using the following primers: SST_1_; CTACTTTGCCGCCTGGTGCTC and TGGCAATGATGAGCACGTAAC, SST_2_; TTGACGGTCATGAGCATC and ACAGACACGGACGAGACATTG, SST_3_; GGCCGCTGTTACCTATCCTTC and GGCACTCCTGAGAACACAACC, SST_4_; CGGAGGCGCTCAGAGAAGAAG and TGGTCTTGGTGAAAGGGACTC and SST_5_; CATGAGTGTTGACCGCTACC and GGCACAGCTATTGGCATAAG, neprilysin; GACTCCCCTGGAGATCAGCCTCTCT and GGTTTTCATCAATAGGCAAAC and for GAPDH; ACCACAGTCCATGCCATCAC and TCCACCACCCTGTTGCTG.

### Osmotic pump administration of an agonist selective to SST_1_ and SST_4_

Two-month-old *App*^*NL-G-F*^ mice were anaesthetized with pentobarbital, the skull exposed, and a hole drilled through the skull with a dental drill. A brain infusion cannula (Alzet cat#8851) connected to an osmotic pump (Alzet cat# 1004) was inserted at bregma −2.5, mediolateral, 2.0 mm and dorsoventral 2.0 mm to administer PBS or 56 μM CH275 (Tocris, cat#2454) dissolved in PBS at 0.11 μl/h which, according to the manufacturer’s instructions (Alzet), would yield a 100 nM concentration in the interstitial fluid (n=12/group). The pumps were changed after 4-6 weeks by surgical intervention. At the age of 6 months, the mice were subjected to behavioral analysis followed by brain isolation. To measure successful delivery of CH275, two weeks administration was performed and then hippocampus was dissected. 30.4 mg hippocampal tissue was homogenized in 500 μl of 10% CH_3_CN-0.1%TFA, the homogenate was sonicated for 10 min and centrifuged at 14,200 × g for 5min. The resulting supernatant was ultrafiltrated with a molecular cut-off weight of 10,000. After centrifugation at 11,000 × g for 1.5 h, the resulting solution (<MW 10,000) was analyzed by reversed phase HPLC. All fractionation steps were performed at 4°C. For quantitation, pure CH275 was injected to the HPLC and used as standard to measure the area under the peak in the resulting diagram. Reversed phase HPLC were carried out using the SHISEIDO CAPCELL PAK C18 (4.6 mm × 250 mm). The mobile phase composed of 0% CH3CN-0.1% TFA for 5 min, linear gradient of 0% to 90% CH3CN-0.1% TFA for 30 min, 90% CH3CN-0.1% TFA for 5 min, and 0% CH3CN-0.1% TFA for 5 min. The flow rate was 0.3 ml/min, and the detection was done at 210 nm.

### I.p. injection of SST_4_ agonist

SST_4_ agonist NNC 26-9100 (Tocris) was dissolved to 90 μM in 95% ethanol and 300 μl injected *i.p.* every second day for two weeks into 6-month-old *App*^*NL-F*^ mice. After two weeks’ administration the brains were removed and analyzed.

### Y-maze

Eighteen-month-old *Sst*_*1*_ KO, *Sst_4_* KO, *Sst_1_xSst_4_* dKO or CH275/PBS-treated *App*^*NL-G-F*^ mice were acclimatized to the behavioral laboratory for two days, and before tests the cages with mice were placed in white noise for one hour. A 12 hr:12 hr light sequence (lights on at 8:00 am) was used. The laboratory was air-conditioned and maintained at a temperature of approximately 22–23 °C and relative humidity of approximately 50–55%. Food and water were freely available except during experimentation. Tests were performed from 9:30 am to 3:30 pm. The Y-maze apparatus (O’Hara) was made of gray plastic and consisted of three compartments (3-cm (width) bottom and 10-cm (width) top, 40 cm (length) and 12 cm (height)) radiating out from the center platform (3 × 3 × 3 cm triangle). Each mouse was placed at the end of one arm facing the wall and was then allowed to explore freely for 5 min. Experiments were performed at a light intensity of 150 lux at the platform. As a measure of memory, the percentage alternation between the visits of the arms of the maze was calculated.

### Morris water maze

Eighteen-month-old *Sst*_*1*_ KO, *Sst_4_* KO and *Sst_1_xSst_4_* dKO mice were transferred to the experimental room the day before the test day. Each mouse was assessed in two training sessions per day. Mice had ad libitum access to food and water (except during the tests). The water temperature was 24 °C and four cues were places around the pool. The mice were allowed to swim for one minute and if they had not reached the platform within one minute, they were placed manually on the hidden platform before being put back in their cage. The mice were trained for five days and the latency time for each mouse strain was analyzed.

### Open field test

CH275/PBS-treated *App*^*NL-G-F*^ mice were transferred to the experimental room at the beginning of the test day and left for one hour in white noise before the start of the test. Mice had ad libitum access to food and water (except during the tests). The mice were placed in the middle of the box and allowed to explore the surface for ten minutes. The amount of time spent in the center region was used as a measure of anxiety.

### Quantitative reverse transcription polymerase chain reaction

Total RNA was extracted from tissues of 13-month-old wildtype C57BL/6J mice (n=3) using RNAiso Plus (Takara Bio). Complementary DNA (cDNA) was synthesized from 1 μg total RNA using ReverTra Ace^®^ qRT-PCR RT Master Mix with gDNA Remover (Toyobo, Osaka) according to the manufacturer’s protocol. The cDNA was used for quantitative PCR with Realtime PCR Master Mix (Toyobo, Osaka) and a TaqMan array using the ABI 7900HT system (Life Technologies). The primer and probe sets for mouse GAPDH (Mm99999915), SST_1_ (Mm00436679) and SST_4_ (Mm00436710) were purchased from Applied Biosystems. GAPDH was used to normalize the quantitative RT-PCR values.

### Statistical analysis

The experimental data were analyzed by Student’s t-test.

